# Intra- and interspecific relationships of alpine ungulates reorganise along a wolf recolonisation gradient

**DOI:** 10.64898/2026.10.12.758551

**Authors:** Charlotte Vanderlocht, Andrea Corradini, Luca Corlatti, Valerio Donini, Francesco Ferretti, Laura Limonciello, Marco Musiani, Federico Ossi, Noemi Squillaci, Laura Viviani, Luca Pedrotti, Heidi C. Hauffe, Francesca Cagnacci

## Abstract

Apex predators are recovering across many ecosystems after decades of decline, yet whether their return can restore ecological interactions disrupted by their loss remains poorly understood. Here, we used the ongoing recolonisation of grey wolves (*Canis lupus*) in the south-eastern Alps as a natural gradient of predator recovery to examine how predation risk and human disturbance shape relationships among red deer (*Cervus elaphus*), roe deer (*Capreolus capreolus*), and Alpine chamois (*Rupicapra rupicapra*). Across a gradient of time since wolf establishment, roe deer increasingly occupied sites without red deer, suggesting predator recovery alters relationships between prey species. Red deer also formed larger groups, consistent with a shift in collective behaviour under increased predation risk. Human disturbance also affected group formation in red deer and chamois. Together, these findings show that prey community structure can vary with wolf recovery, even in human-dominated landscapes, through altered associations among species and grouping within species.

## Introduction

Humans are reshaping the natural world at an unprecedented pace^1^, accelerating biodiversity loss^2^ with consequences that extend beyond species extinctions to the disruption and reorganisation of ecological interactions^3^. Among the most pervasive human impacts is the widespread loss of apex predators, which has weakened their regulatory influence on prey populations and behaviour – a process known as trophic downgrading^4^. Predator loss can therefore alter not only prey abundance and behaviour, but also the interactions among species, with effects that can cascade through ecological communities^5,6^. Such changes can ultimately alter community structure, energy flow, and ecosystem functioning^7^.

Yet the ecological consequences of trophic downgrading may not be irreversible^8^. Large carnivore populations are recovering across parts of their historical ranges in Europe and North America, decades to centuries after local extirpation^9,10^. The ecosystems to which these apex predators return, however, have been profoundly transformed by human activities^11,12^, with degraded and fragmented habitats, altered species assemblages, and persistent human disturbance potentially shaping species interactions in ways that differ from the past. Therefore, the return of apex predators provides a large-scale natural experiment to examine whether, and to what extent, complex ecological interactions can be restored in human-modified ecosystems. Here, we address this challenge by leveraging the ongoing recolonisation of grey wolves (*Canis lupus*) in the southern Alps to evaluate its impact on the behaviour of an ungulate guild.

A growing body of literature shows that prey may adjust their behaviour following the return of their predators^13^, including changes in vigilance^14^, daily activity patterns^15^, and habitat or resource use^16^. While these studies provide crucial insights, they have primarily focused on single-species responses. Ecological communities, by contrast, are structured by interactions among coexisting organisms within shared environments^17^, yet how predator recovery reorganises interactions *at the community level* remains poorly understood. Such community-level shifts are simultaneously shaped by interspecific and intraspecific associations, which together determine how species and individuals are distributed across space and time, and shape the interactions through which communities assemble and persist across heterogeneous landscapes.

Large herbivore communities provide some of the most striking examples of such structured coexistence. In African savannas, for example, more than 20 ungulate species can occur sympatrically^18^, forming large and diverse mixed-species aggregations that comprise dozens to hundreds of individuals. While such coexistence has traditionally been explained by resource partitioning^19,20^ and resource facilitation^21,22^, behavioural processes also play a central role. These processes span from direct interference competition and avoidance behaviour^23^ to indirect interactions mediated by shared threats^24^, especially shared predation risk^25^.

Among these indirect interactions, apparent competition can arise when species indirectly affect one another via predator-risk-mediated responses^26^. These negative indirect interactions can cause species that differ in vulnerability to predation to use the landscape differently, where more vulnerable species avoid areas associated with less vulnerable species^27,28^. Such dynamics have been described in systems involving caribou (*Rangifer tarandus*), moose (*Alces alces*), and wolves (*Canis lupus*). Conversely, shared predation risk can also generate positive interspecific associations. Mixed-species co-occurrence may provide antipredator benefits such as increased vigilance, as observed among African buffalo (*Syncerus caffer*), Burchell’s zebra (*Equus quagga*), and wildebeest (*Connochaetes taurinus*)^29^; access to social information, as documented in mixed-species bird communities^30,31^; or reduced species-specific or individual risk through dilution effects^32^. These benefits may be particularly pronounced when species differ in predator detection abilities, allowing one species to benefit from the vigilance or early-warning cues of another (the protector-species hypothesis), as proposed for associations between black skimmers (*Rhynchops niger*) and gull-billed terns (*Sterna nilotica*)^33^. Together, these behavioural processes can shape complex patterns of spatiotemporal co-occurrence that cannot be predicted from resource use alone^25,34,35^, but instead emerge from trade-offs between antipredator benefits and foraging constraints in large herbivores^36^.

Critically, similar trade-offs also structure intraspecific grouping across a wide range of social and gregarious animals, including fishes, birds, and mammals. Individuals can balance the antipredator benefits of collective vigilance (the “many-eyes” effect^32^), reduced individual risk when danger is shared among group members (the “dilution of risk” effects^37^), and positional advantages (the “selfish herd” effect^38^) against costs including competition^39^, increased detectability^40^, and social distraction^41^.

These inter- and intra-specific associations among prey species may be expected as apex predators recover. In human-dominated landscapes, however, prey behavioural responses also depend on anthropogenic pressures. Humans can act as ‘super-predators’^42^, inducing widespread behavioural responses in wildlife^43^. Non-lethal human disturbance can elicit behavioural responses analogous to those triggered by natural predators^44^, with effects that vary with the predictability of disturbance^45^ and its spatial form, ranging from static infrastructure to recreational human use^46,47^.

Here we investigate how predator recovery and anthropogenic disturbance jointly structure behavioural organisation within an Alpine large herbivore community, including red deer (*Cervus elaphus*), roe deer (*Capreolus capreolus*), and Alpine chamois (*Rupicapra rupicapra rupicapra*). In the south-eastern Alps, these species have undergone substantial population recoveries following a mid-20th century bottleneck^48^. During summer, their ranges overlap extensively due to altitudinal migration^49–51^ and locally high resource availability^52^, creating strong potential for co-occurrence and behavioural interactions. This ecosystem is exposed to ongoing wolf recolonisation^53^ (Fig.1A) and increasing non-lethal human disturbance, especially tourism^48^ and outdoor recreation, particularly during the summer^54^. Wolf predation risk has re-emerged and varied spatially and temporally following more than a century of absence^55–57^. To capture this gradient of time since wolf recolonisation, we used three consecutive years of summer camera-trap data along the recolonisation front (Fig.1B), leveraging both spatial differences among sites and temporal variation in time since wolf establishment within sites, while accounting for environmental covariates, to test whether inter- and intraspecific prey associations systematically tracked predator recovery. We quantified interspecific co-occurrence and intraspecific grouping patterns in Alpine ungulates and evaluated how these varied along gradients of wolf recolonisation (Fig. 1A-B), human disturbance (Fig.1C), and environmental conditions such as tree cover density and terrain steepness, in relation to species-specific traits (Fig.1D).

**Figure 1.**
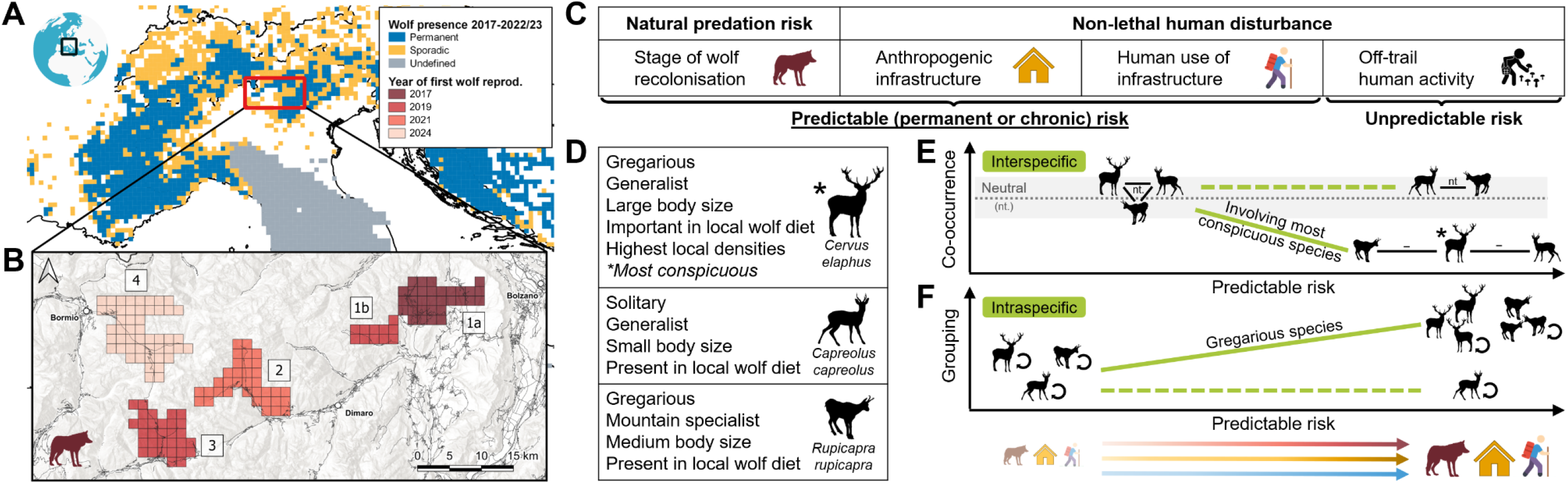
Study design and conceptual predictions. (**A**) Wolf distribution across the Alps from 2017–2022/23^105^. Permanent, occasional, and undefined (i.e. wolf presence is established but its status remains unclear) occurrences are shown in dark blue, yellow, and grey, respectively. (**B**) The four study areas span a temporal gradient since wolf recolonisation(shades of red): from right to left: 1a – Upper Non Valley (2017 – first year of wolf reproduction); 1b – Bresimo Valley (2019); 2 – Peio Valley (2021); 3 – Tonale Pass (2019); 4 – Valfurva (2024). (**C**) Risk factors examined in the study included natural predation risk, captured through quasi-experimental variation in time since wolf recolonisation across study areas, and non-lethal human disturbance. Human disturbance was characterised using three complementary proxies differing in predictability and spatial structure: permanent anthropogenic infrastructure, predictable human use of infrastructure, and unpredictable off-trail human activity. (**D**) The alpine ungulate community and the main species-specific traits expected to influence behavioural responses to risk. (**E**-**F**) Conceptual predictions for inter- and intraspecific ungulate responses to increasing predictable risk, depending on species-specific traits. Light green lines indicate expected responses; solid lines denote strong modulation by predictable risk, whereas dashed lines indicate weak or no modulation. Risk gradients shown below panels E-F represent increasing predation risk associated with wolf recolonisation (red gradient), and increasing human disturbance from permanent infrastructure (yellow gradient) and predictable use of infrastructure (blue gradient). (**E**) Predicted interspecific associations, expressed as the intensity of co-occurrence between species pairs. The grey dotted line and shaded confidence interval represent neutral co-occurrence (i.e. not significantly different from random expectation, also indicated as “nt.”); values below neutral co-occurrence indicate negative co-occurrence or interspecific avoidance (also indicated as “–”). (**F**) Predicted intraspecific associations, expressed as grouping intensity within species, with increasing values corresponding to larger group sizes.

First, while accounting for the habitat use of each species, we assessed whether pairwise co-occurrence among the three ungulate species deviates from random expectations, and whether such patterns are associated with wolf recolonisation and/or anthropogenic disturbance. We used multi-species occupancy modelling^58^ to test whether species associations are structured beyond expectations from independent habitat use alone. In the absence of external risk, we expected neutral co-occurrence among sympatric ungulates — that is, not more nor less frequent than expected by chance (Fig.1E), as niche segregation likely occurs at finer resource scales^59^. Under increasing predictable risk from apex predator recovery and human disturbance, we predicted that avoidance may emerge among ungulates, consistent with risk-mediated apparent competition. Specifically, we expected this to occur for ungulate pairs involving red deer, which may be more conspicuous and thus more easily detected by wolves, because of their larger body size, gregariousness, and high local density (locally up to 30 ind/km²; Fig.1E).

Second, we examined how the same set of drivers influences intraspecific grouping behaviour, testing whether group size in each species varies with time since wolf recolonisation, human disturbance, and environmental context, including vegetation cover and topography. We modelled intraspecific group size as observed on independent camera-trap events. Under increasing predictable risk from apex predator recovery and human disturbance, we predicted increased group size in gregarious species (red deer and chamois; Fig.1F), reflecting dilution and collective vigilance effects. Although roe deer are capable of context-dependent grouping in winter and early spring^60^, we did not expect groups to form in summer, when the species is more solitary, nor that it would be affected by risk (Fig.1F). We also did not expect unpredictable human activity to influence interspecific or intraspecific associations at the scale under consideration.

## Results

Camera-trap deployments throughout three consecutive summers across 355.5 km² resulted in 57,091 effective camera-days, yielding a total of 6,231 independent sightings of red deer, 2,608 of roe deer, and 933 of chamois.

### Interspecific co-occurrence

Species occupancy depended on environmental conditions but not on the presence of heterospecifics. After accounting for species-specific habitat use, pairwise co-occurrence varied with wolf recolonisation and terrain steepness.

Correcting for imperfect detection (Supporting information Table S1), the estimated probability that a site was occupied was similar for red deer and roe deer across the study area and sampling period, with mean occupancies [95% CI] of 0.85 [0.64–0.93] and 0.90 [0.83–0.95], respectively (Fig.2). Chamois, on the other hand, had an occupancy of 0.57 [0.40–0.72] (Fig.2).

**Figure 2.**
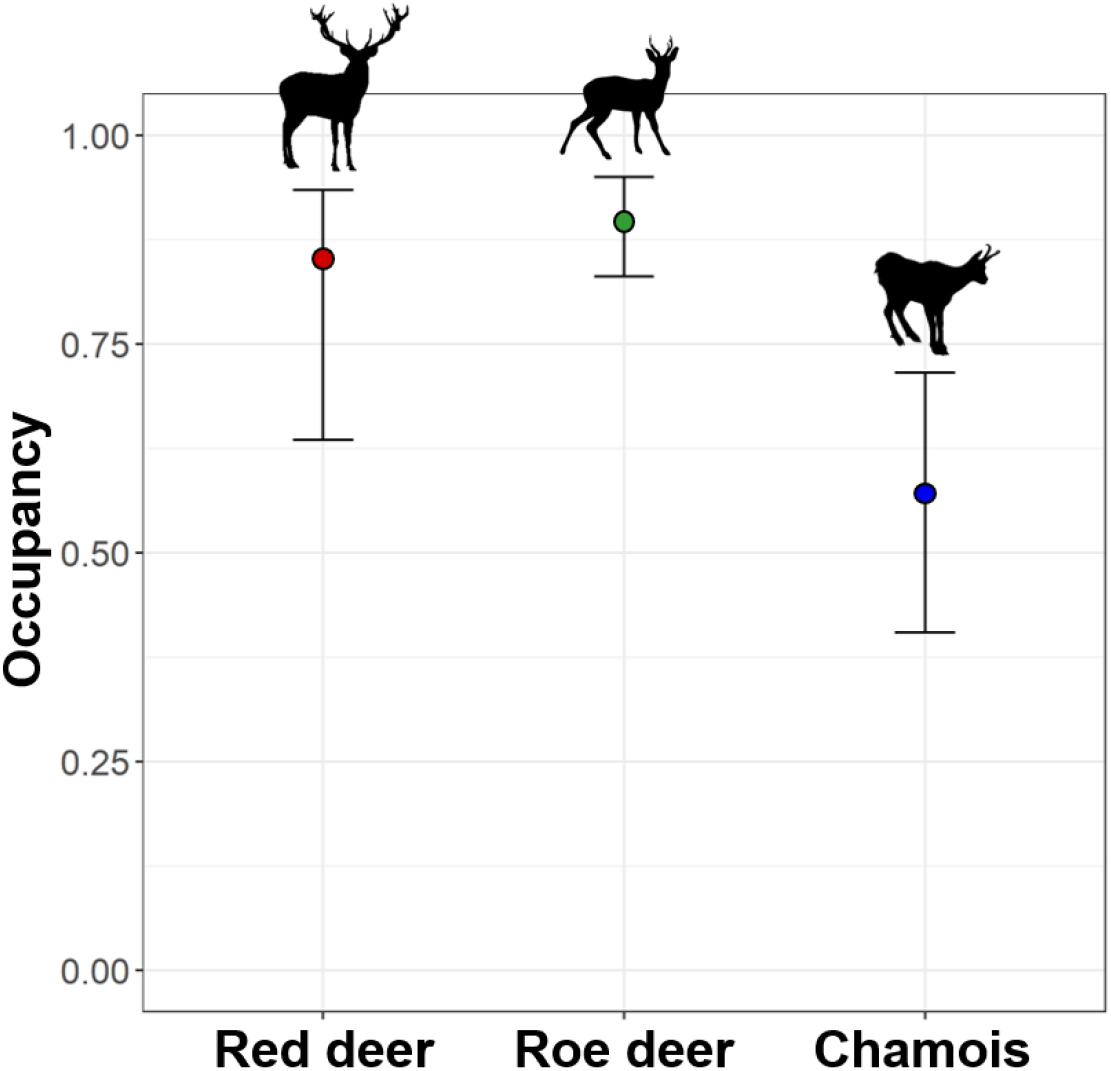
Occupancy of red deer (red), roe deer (green) and chamois (blue) throughout the study area and period. Coloured dots show mean probability of occupancy and lines the 95% confidence intervals. Occupancy probabilities were estimated by setting all covariates to their mean values.

In relation to environmental and risk variables, chamois occupancy probability increased with terrain slope (β_slope_ = 0.84 ± 0.39), while roe deer occupancy was higher in densely forested areas (β_tree cover density_ = 1.29 ± 0.41) and in proximity to human settlements (β_distance to nearest settlement_ = -1.04 ± 0.43). Occupancy of roe deer was also positively associated with time since wolf recolonisation (β_time since first wolf reproduction_ = 0.72 ± 0.36). Occupancy of red deer decreased with predictable recreational human activity (β_density of cumulated outdoor activity index_ = -0.66 ± 0.30).

The overall occupancy of each species was unaffected by the presence or absence of other species (Supporting information Fig.S1).

When considering the effects of environmental conditions, anthropogenic pressures, and wolf establishment on pairwise co-occurrence patterns, we found that co-occurrence between red deer and roe deer was influenced by time since wolf establishment (β_time since first wolf reproduction_ = -0.92 ± 0.37), but with asymmetric effects in magnitude between the two cervids. As time since first wolf reproduction increased, roe deer occupancy in the absence of red deer increased more than threefold (from 0.19 to 0.77 occupancy probability over a decade of gradual wolf recolonisation; Fig.3A) and red deer occupancy in the presence of roe deer decreased from 0.98 to 0.87 (Fig.3B). Co-occurrence between red deer and chamois was affected by terrain slope (β_slope_ = -0.85 ± 0.40), with a negative association strengthening on steeper slopes: chamois occupancy in the absence of red deer increased strongly with slope (more than 21-fold from 0.03 to 0.72 occupancy probability across slopes ranging from 0° to 45°; Fig.3C), while slope effects on red deer occupancy were weak and uncertain, with overlapping confidence intervals across conditions defined by chamois presence (Fig.3D). Model details are reported in Supporting information Table S2.

**Figure 3.**
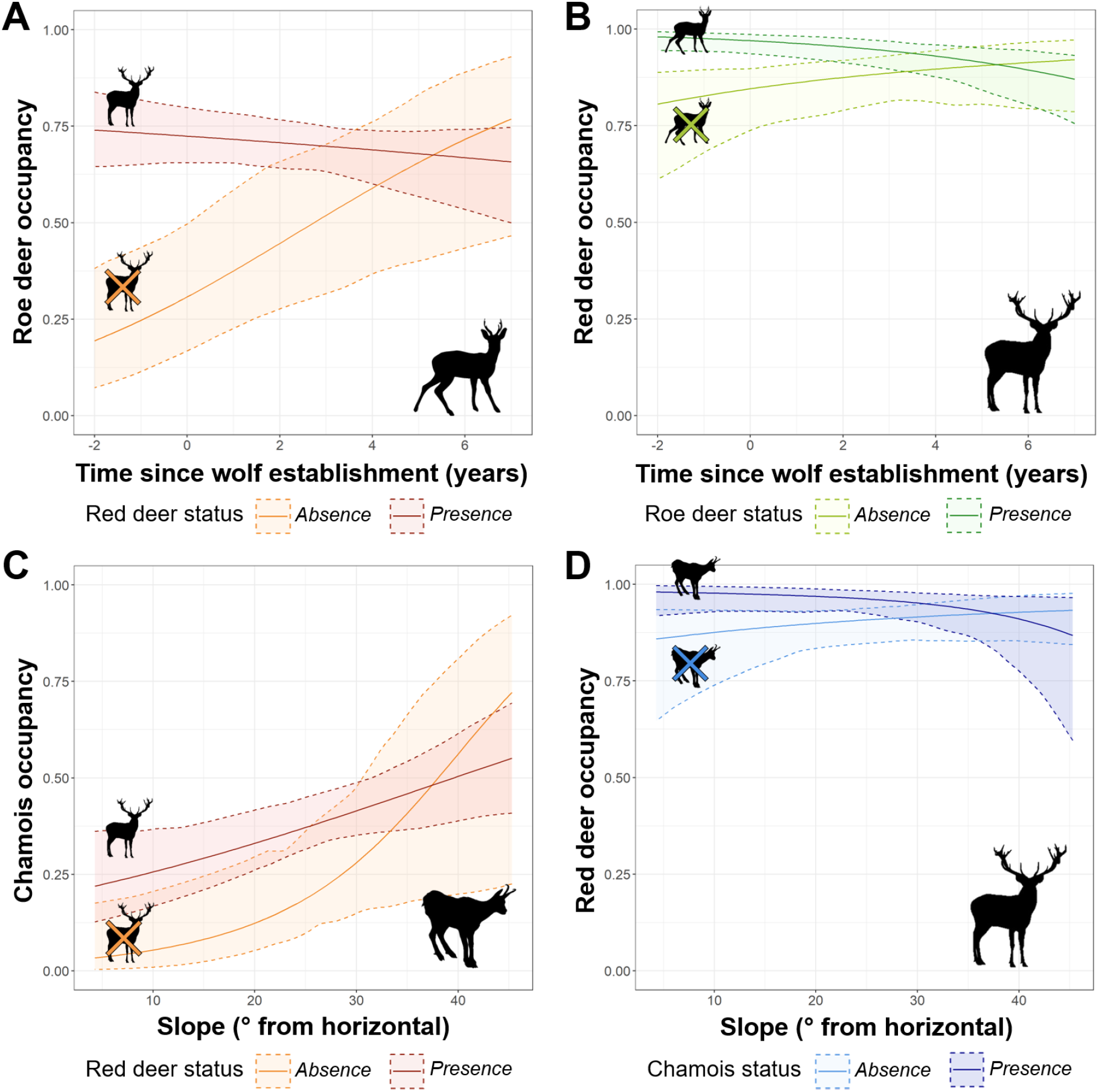
Occupancy of (**A**) roe deer, (**B,D**) red deer, and **(C)** chamois as a function of (**A-B**) time since wolf establishment (in years, with 0 for the year of first wolf reproduction and negative values for years prior to wolf reproduction) and as a function of (**C-D**) slope, in each other’s presence or absence (red deer presence in red; red deer absence in orange; roe deer presence in dark green; roe deer absence in light green; chamois presence in dark blue; chamois absence in light blue). Solid lines show mean probability of occupancy and dashed lines the 95% confidence intervals. Occupancy probabilities were estimated by setting all other covariates to their mean values.

### Intraspecific group size

Red deer and chamois group size was associated with gradient of wolf recolonisation, anthropogenic disturbance, and environmental conditions, whereas roe deer group size remained unchanged.

Specifically, estimated red deer group size ranged from 1 to 39 individuals (with a mean of 1.74 and median of 1). Group size was smaller on steep slopes (β_slope_ = -0.06 ± 0.01; Fig.4A), and larger near human settlements (β_distance to nearest settlement_ = -0.06 ± 0.01; Fig.4C).

**Figure 4.**
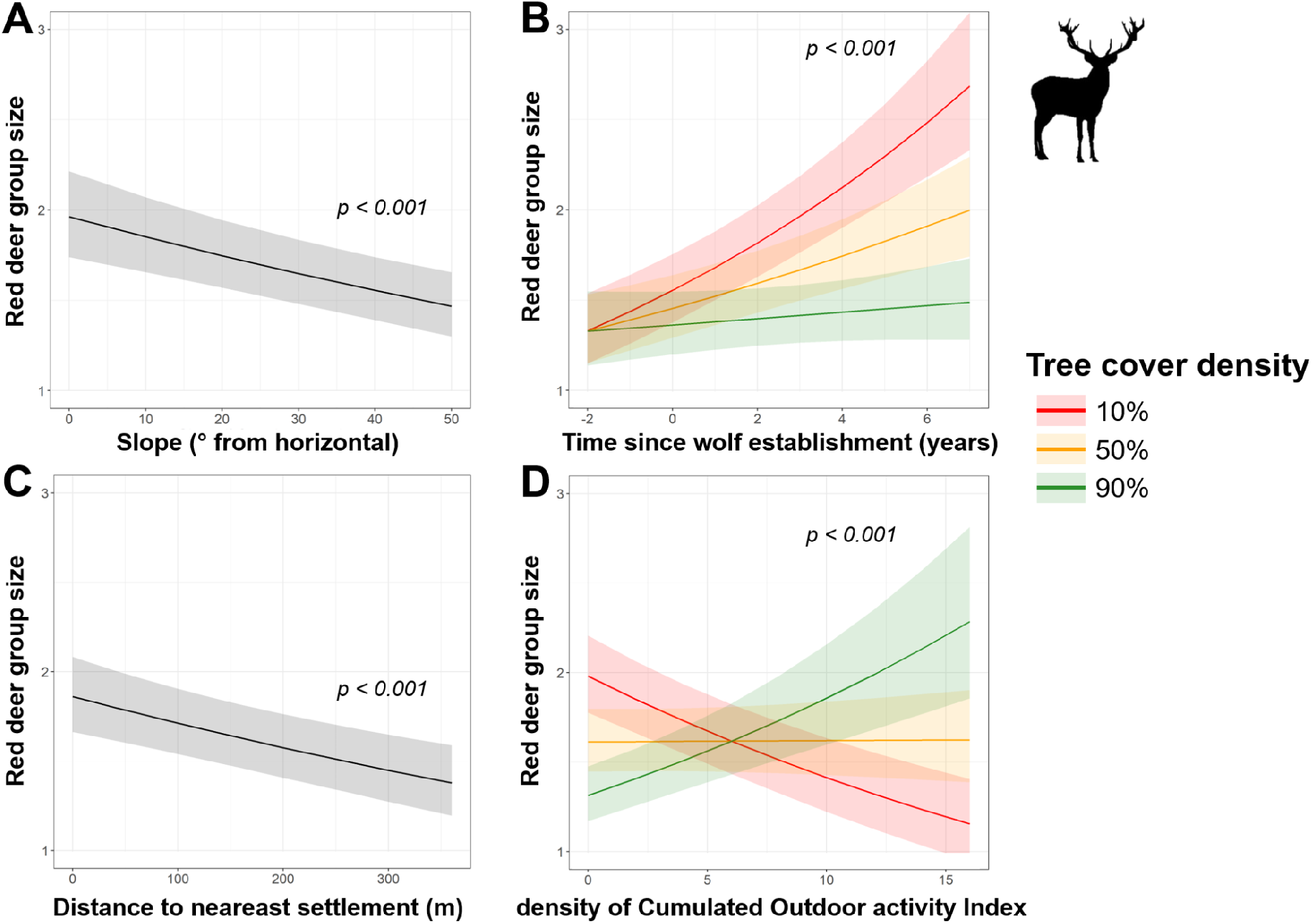
Predictions of red deer group size as a function of (**A**) slope, (**B**) time since wolf establishment (in years, with 0 for the year of first wolf reproduction and negative values for years prior to wolf reproduction) in interaction with tree cover density, (**C**) distance to nearest settlement, and (**D**) density of Cumulated Outdoor activity Index in interaction with tree cover density. Lines with shaded areas represent the model prediction (mean effect with 95% CI), and tree cover density is plotted for 10% (red), 50% (yellow), and 90% (green).

Red deer group size increased with time since wolf establishment, especially so in more open areas (β_time since first wolf reproduction x Tree cover density_ = -0.08 ± 0.01; Fig.4B). With increasing recreational space use by humans, red deer group size increased in forested areas, but decreased in open areas (β_density of Cumulated Outdoor activity Index x Tree cover density_ = 0.09 ± 0.01; Fig.4D).

Chamois group size ranged between 1 and 15 individuals (with a mean of 1.47 and median of 1). As for red deer, group size was also smaller on steep slopes (β_slope_ = -0.12 ± 0.03; Fig.5A). With increasing proximity to human settlements, chamois group size increased in forests, but not in open areas (β_distance to nearest settlement x Tree cover density_ = -0.07 ± 0.04; Fig.5B). With increasing recreational use of space by humans, chamois group size decreased in forested ^areas, but not in open areas (β^_density of Cumulated Outdoor activity Index x Tree cover density_ ^= -0.11 ± 0.04;^ Fig.5C).

**Figure 5.**
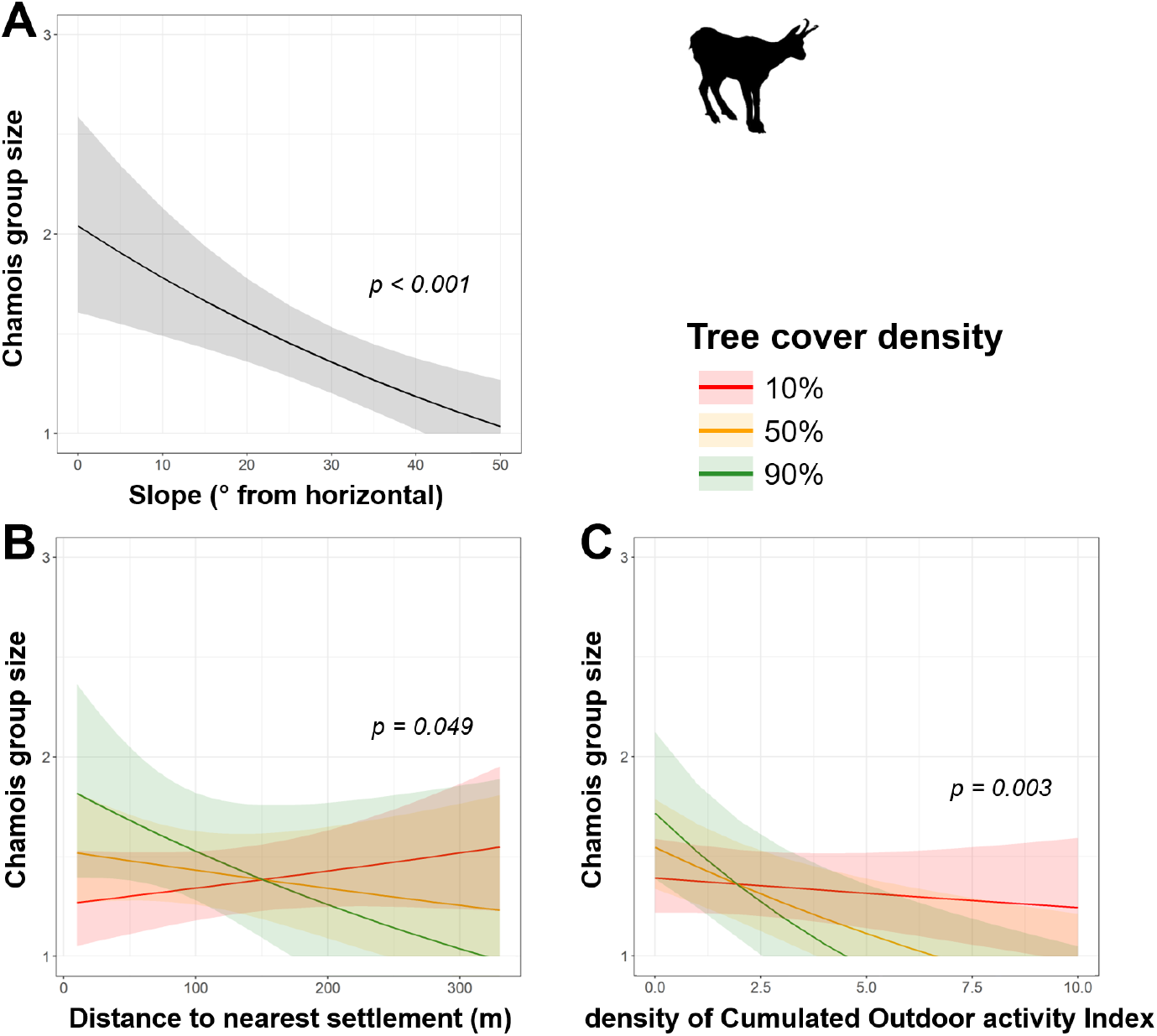
Predictions of chamois group size as a function of (**A**) slope, (**B**) distance from nearest settlement in interaction with tree cover density, and (**C**) density Cumulated Outdoor activity Index in interaction with tree cover density. Lines with shaded areas represent the model prediction (mean effect with 95% CI), and tree cover density is plotted for 10% (red), 50% (yellow), and 90% (green).

Roe deer group size ranged between 1 and 6 individuals (with a mean of 1.18 and median of 1), and was not affected by any environmental or risk variable measured here. Model details are reported in Supporting information Table S3.

## Discussion

Our results show that the natural return of the grey wolf to a human-dominated alpine landscape is associated with a complex reorganisation of the large herbivore prey community structure, with shifts in both interspecific associations and intraspecific grouping. As time since wolf establishment increased, roe deer were more likely to occupy sites where red deer were absent, suggesting an avoidance mechanism consistent with predator-mediated apparent competition, whereby species differed in their tolerance of shared predation risk (i.e. wolf predation; see below). Red deer formed increasingly larger intraspecific groups as time since wolf establishment increased, particularly in open habitats, suggesting an emerging pattern of ‘risk-sensitive grouping’, whereby aggregation provides antipredator benefits through dilution and collective vigilance^32,40,61,62^. We found no evidence that anthropogenic pressures (that is, distance to human settlements, density of cumulated outdoor activity index, and likelihood of off-trail human detection) influenced interspecific co-occurrence of ungulates. However, permanent infrastructure (distance to human settlements) and predictable on-trail human activity (density of cumulated outdoor activity index) did shape intraspecific grouping patterns in red deer and chamois, depending on habitat openness. Patterns of co-occurrence between red deer and chamois, as well as intraspecific grouping behaviour for both species, were shaped by terrain slope: although both species individually occupied steep areas, they formed smaller groups and reduced interspecific aggregation on such terrain, highlighting shared environmental constraints that influence both inter- and intraspecific associations.

### Roe deer and red deer co-occur less after the wolf’s return: predator-mediated apparent competition at its onset?

Patterns of co-occurrence between roe deer and red deer suggested asymmetric behavioural responses to shared predation risk along the wolf recolonisation gradient. Because our models accounted for species-specific responses to environmental and anthropogenic variables, this pattern is unlikely to be simply explained by shared or diverging habitat preferences. We found no evidence that the biophysical environment changed during the study, with the exception of wolf presence. Finally, in our study the two species did not avoid each other in the absence of wolves, which is consistent with limited niche overlap in terms of dietary preferences^63–65^. Instead, our results suggest differing costs and benefits of co-occurrence under shared predation risk^25,26^. Overall, these patterns suggest a stronger spatial decoupling between roe deer and red deer than is typically reported in European systems where wolves are or were absent, where the two cervids often coexist with substantial spatial and habitat use overlap despite fine-scale niche segregation^66,67^.

While our model focussed on pairwise co-occurrence, predicted model outputs revealed marked asymmetry in species responses, with roe deer showing a stronger increase in occupancy in the absence of red deer under increasing wolf establishment, whereas red deer exhibited only a weaker reduction in occupancy in the presence of roe deer. This pattern suggests that roe deer are more sensitive to changes in heterospecific context under elevated predation risk. Given their smaller body size, more solitary behaviour (see absence of grouping responses in Supporting information Table S3), and generally lower conspicuousness, roe deer may lose their relative crypsis when associating with the larger, more gregarious red deer, potentially increasing exposure to wolf predation.

More broadly, our results highlight that coexistence is shaped not only by competition or facilitation, but also by behavioural trade-offs under shared risk^23,68^. These responses align with behavioural mechanisms -underlying and/or preceding predator-mediated apparent competition, where asymmetrical vulnerability leads one species to avoid another to reduce risk exposure^25,28^. While apparent competition is often considered in demographic terms^34,69^, it can also unfold behaviourally through avoidance, altered space use^28^, or shifts in activity patterns, including in the early stages of predator colonisation. In the near future, at a more advanced stage of wolf recolonisation, demographic analysis will be needed to assess whether the patterns observed here will translate into population consequences. As DeCesare et al.^34^ underscore, such interactions can have significant demographic consequences later on, particularly as predator communities reassemble. Finally, while our 48-hour temporal resolution of co-occurrence data offers meaningful insights, finer temporal analyses are needed to determine whether avoidance or attraction patterns occur in real time or with delays: this will be critical for understanding the mechanisms underlying species interactions^27,28^.

### Red deer adjust grouping behaviour in exposed habitats as wolves recolonise the Alps

Red deer formed increasingly larger intraspecific groups with time since wolf establishment, particularly in open habitats. This behaviour aligns with classic antipredator strategies, whereby group formation dilutes individual risk and enhances collective detection in environments where predation risk is elevated^38,70^. Similar context-dependent grouping responses have been observed in elk (*Cervus canadensis*), which formed either smaller groups to reduce encounter rates or larger ones to dilute risk, depending on habitat and individual life history^71^. Open habitats, while offering foraging benefits, may increase exposure to predators, thereby incentivising social aggregation. Indeed, musk-oxen formed larger groups in high-wolf-risk areas^70^, and caribou were more social in open habitat and during nocturnal periods when risk is higher^72^. Our findings echo these patterns, suggesting that grouping can help red deer to mitigate the costs of occupying productive yet risky open areas under growing wolf pressure. While group size can also be influenced by local density^60,71,73^, our model structure accounted for study area-level differences via random effects. Nevertheless, future work would benefit from explicitly incorporating local density, sex-ratio, as well as sex and age within groups, to disentangle behavioural from demographic drivers of grouping dynamics.

### Gregarious alpine ungulates adjust grouping behaviour to non-lethal human disturbance

Both red deer and chamois adjusted group size along gradients of predictable forms of non-lethal human disturbance, namely proximity to anthropogenic infrastructure and the intensity of human use quantified through a cumulative outdoor recreation index^47^. Predictability, or chronicity, of risk is a key driver of proactive antipredator responses such as changes in group size^74^, whereas less predictable disturbances, such as direct off-trail human activity, are more likely to elicit reactive behavioural responses^75,76^.

Red deer formed larger groups near human settlements, suggesting that predictable human presence favours collective vigilance as a strategy to mitigate perceived risk. Vicinity to human settlements may also correlate with greater forage availability, such as managed meadows, potentially attracting deer to otherwise riskier areas, where collective vigilance helps offset the costs of increased human exposure. Responses to outdoor recreation, however, depended strongly on habitat structure. Red deer formed larger groups in disturbed forest but smaller groups in disturbed open habitats, highlighting how vegetation cover modulates perceived risk and opportunities for concealment^76–78^. In forested environments, increased grouping could facilitate collective vigilance where visibility is limited^79^, whereas in open habitats, where animals are already highly conspicuous, the costs of increased detectability may outweigh the antipredator benefits of aggregation. Similarly to red deer with human disturbance, chamois formed larger groups near human settlements embedded within forested landscapes, suggesting that structurally protective habitats similarly favour collective vigilance under predictable human presence. However, in forests subject to high recreational use, chamois instead formed smaller groups, potentially prioritising reduced detectability and crypsis over the benefits of collective vigilance.

Overall, our findings highlight that behavioural responses to non-lethal human disturbance are highly nuanced and context-dependent^80^. In some settings, gregarious ungulates responded in ways consistent with classical antipredator theory^44^, increasing group size to enhance collective vigilance or dilution of risk. In others, however, they appeared to minimise exposure by reducing detectability, suggesting that responses to humans cannot always be interpreted solely through the lens of predation risk. Rather, behavioural responses to human presence may depend on how disturbance is experienced, including its predictability and habitat structure. Future studies could further disentangle the behavioural trade-offs underlying social organisation in increasingly human-dominated landscapes, for example, by integrating the sex-age structure.

### Inter- and intraspecific aggregation in red deer and chamois is less observed on steep terrain

The importance of slope in occupancy patterns in red deer and chamois offers an example of how the heterogenous environmental structure of alpine environments can shape both inter- and intraspecific spatial dynamics. Chamois showed a clear association with steeper slopes, and red deer a positive, but marginal, association as well. However, the two species co-occurred less on steeper slopes, suggesting spatial segregation within commonly used terrain. Interestingly, a comparable pattern was evident within species: both red deer and chamois formed smaller intraspecific groups in steeper areas, indicating that individual occurrence became increasingly decoupled from group formation with increasing slope.

Steep rocks are known refuge areas for mountain ungulates^81–83^, thus our findings may indicate that steep terrain selection constitutes a distinct antipredator strategy from grouping. Furthermore, steeper slopes are often characterised by rockier substrates and sparser vegetation, which can reduce foraging opportunities and make aggregation less energetically favourable^84^. From a methodological perspective, detectability of camera-trap data may vary with terrain complexity despite standardised deployment protocols^85–87^, a limitation common to many camera-trap studies. Future analyses incorporating explicit detection submodels or complementary data sources (e.g., direct observations or drone-based surveys) could further refine our understanding of aggregation across complex terrain. Understanding how species navigate these structurally complex habitats, and how behaviour constrains or facilitates coexistence within them, will be critical to anticipating the spatial reorganisation of herbivore communities under several types of risk, resource availability, and climate change in mountainous environments.

## Conclusions

Across many landscapes, ungulates are becoming increasingly exposed to recovering large carnivore populations alongside persistent human activity^9,10,12,88^. Our study demonstrates that these concurrent pressures are jointly associated with the restructuring of behavioural relationships across and within species, revealing an overlooked dimension of community reassembly during predator recovery. More broadly, our results illustrate the value of ongoing predator recolonisation as a large-scale natural experiment. Because recolonisation unfolds progressively across landscapes, it provides a unique opportunity to observe how ecological interactions emerge, reorganise, and potentially stabilise over the course of only a few years. Following these systems through time will allow us to move beyond documenting behavioural responses in individual species towards understanding how foundational ecological relationships – including coexistence, indirect interactions, and the organisation of communities – are re-established in ecosystems recovering from trophic downgrading. Such long-term perspectives will be essential for linking behavioural reorganisation to demographic processes, species coexistence, and ecosystem functioning in increasingly human-dominated landscapes.

## Methods

### Study areas and study design

The study area was located in the south-eastern Alps (Fig.1A), in the Italian regions of Lombardy and Trentino-Alto Adige/Südtirol, including the Stelvio National Park (IUCN category II). The landscape is characterised by rugged mountainous terrain (900–3,905 m a.s.l.), with alpine pastures at higher elevations, and coniferous forests with managed meadows at lower elevations. We exploited ongoing natural wolf recovery to implement a quasi-experimental design based on a spatial gradient of wolf establishment timing across four study areas (total area of 355.5 km²; mean study area size: 88.9 km² [min: 69.75; max: 112.5]; see also^15^; Fig.1B). These areas were selected to be broadly comparable in landscape structure while differing in the timing of reproductive wolf pair establishment (2017, 2019, 2021 and 2024), providing a proxy gradient for increasing duration of wolf presence.

### Standardised camera-trapping

The total sampling area consisted of a total of 158 gridcells (1.5 x 1.5 km each), with one camera-trap placed randomly in each cell, following a standardised protocol to maximise homogeneity of detection rates and ranges (50 cm height, north-facing, unbaited; Supporting information). The sampling period spanned from May to October, but for the purpose of this study, we focused on events from July and August of 2022–2024. This period was selected to minimise the influence of seasonal movements, as spring and fall migration occur before and after the study period, respectively, thereby better approximating population closure in a natural system^89^. Additionally, summer conditions reduce the influence of winter constraints, such as snow cover and resource scarcity, on habitat use, allowing observed patterns of overlap, segregation, or grouping to better reflect interactions and habitat preferences.

During this period, motion-triggered cameras operated with minimal trigger delays (1–5 s, depending on the model). Field sites were regularly visited to replace batteries and memory cards. All camera-trap media files were annotated using Timelapse^90^ (https://timelapse.ucalgary.ca), producing a dataset that included all detected species at each camera-trap location, the number of individuals per sex and age class (juvenile, yearling, adult, or unknown age class, and female, male, or unknown sex) for each media file, and the corresponding timestamp.

### Environmental and risk variables

To test for the effects of environmental conditions on inter-specific co-occurrence and intra-specific group size, we used variables known to affect ungulate space use and behaviour in Alpine environments: tree cover density^91^, and slope and northness, derived from the available digital elevation model^92^ (Supporting information).

We used the year of first local wolf reproduction as a temporal reference point during recolonisation to capture increasing duration of prey exposure to established wolf presence. This milestone marks the establishment of a stable territory and is associated with increased predation pressure^93^, while also being consistently detected across study sites using opportunistic camera-trapping and genetic monitoring^94,95^. Because wolves may be present prior to reproduction as transient individuals or non-reproductive pairs^96^, ungulates can experience and respond to wolves throughout earlier stages of recolonisation. Time was therefore modelled as a continuous variable spanning three years before to six years after first reproduction, with year 0 indicating its onset and negative values corresponding to pre-reproduction years (-2 to +7 years; Fig1B).

As a proxy for the impact of permanent infrastructure, we used the nearest distance to human settlements^97^. As a proxy for predictable human use of trail infrastructure, we estimated the density of Cumulated Outdoor activity Index (dCOI) based on human recreational mobility data^47^. As a proxy for unpredictable off-trail human activity in the landscape, we calculated the likelihood of detecting a human at the randomly placed camera-trap by counting the number of days with human sightings divided by the number of active camera-trap days. A priori correlation checks confirmed that all these covariates were not correlated (ρ ≤ |0.4|).

### Interspecific co-occurrence modelling

We arranged wild ungulate camera-trap detections into detection matrices with rows representing site-by-year combinations (*i*) and columns corresponding to 48-hour sampling occasions (*j*), applied consistently across all sites and years. This resolution was chosen to capture meaningful behavioural responses while allowing sufficient detection power.

To model detection/non-detection data for the three focal species, we applied a multi-species occupancy framework^58^, which extends the single-species model introduced by MacKenzie et al.^98^ to account for potential co-occurrence patterns among interacting species. The model estimates a latent occupancy state *(Z_i,s_)* for each species *s = 1,…,S* at each site-by-year *i = 1,…,I* where *Z_i_* is a vector of binary values (1 = present, 0 = absent, NA = non-functional camera-trap) describing the presence–absence status across species at site *i*. These latent occupancy states were modelled as a multivariate Bernoulli distribution, *Z_i_* ∼ *MVB(ψ_i_)*, with *ψ_i_*representing the vector of probabilities for each of the possible species combinations (k), with *Σ_k_ ψ_ik_ = 1*. We modelled these probabilities on the logit scale as natural parameters (*f*), capturing the log-odds of occupancy. We focused on pairwise interactions only (no three-way interaction), resulting in a total of six natural parameters per site i: three for single-species occupancy (f*_Red deer,i_*, f*_Roe deer,i_*, f*_Chamois,i_*) and three for pairwise species co-occurrence (*f_Red deer:Roe deer,i_, f_Roe deer:Chamois,i_, f_Chamois:Red deer,i_*).

We used the occuMulti() function in the *unmarked* R package^99^, to model all natural parameters as linear functions of tree cover density, slope, northness, distance to nearest settlement, dCOI, likelihood of human detection, and time since wolf establishment. This allowed us to assess their effects on co-occurrence patterns while accounting for their role in single species occupancy. Finally, observed detection data (*y_sij_*) were linked to the true latent occupancy states through a Bernoulli observation model: *y_sij_* ∼ *Bernoulli(z_si_ p_sij_)*, where *p_sij_* represents the detection probability for species *s*, site *i*, and sampling occasion *j*, assumed to be independent across species and modelled as a function of tree cover density, northness, slope and camera-trap model.

### Intraspecific group size modelling

Following Vanderlocht et al.^15^, we applied a data-driven maximum likelihood approach to determine species-specific time-to-independence thresholds (red deer, roe deer, chamois). In brief, we used the bout-ending criterion^100^, fitting a mixture of two Poisson processes to time intervals between camera-trap sightings, assuming a ‘fast’ process for closely triggered, non-independent images, and a ‘slow’ process for truly independent events at the camera-trap site. We determined the ‘time-to-independence criterion’ as the most likely minimum time period needed to distinguish two independent camera-trap events^101^. This approach enabled us to ensure data independence in the statistical analysis, while remaining sensitive to the different biologies of the species (e.g. sociality, speed of movement, and recursion frequency). We estimated that consecutive images of red deer, roe deer, and chamois were most likely independent if separated by at least about 19, 14, and 21 minutes, respectively (Supporting information Fig.S2).

To determine group size per independent event, we applied a conservative approach: for each independent event, we summed the maximum number of individuals observed per sex-and-age category across all associated images. This method minimised the risk of overestimating group size due to repeated images of the same individuals. However, it may have led to underestimation, as unmarked ungulates could not be individually identified across frames.

For each species separately, we investigated how group size varies with environmental and risk variables, using a generalised linear mixed model with a negative binomial error distribution and a log family (*glmmTMB* R package^102^). Specifically, we modelled the group size of each independent camera-trap event as a function of tree cover density, slope, northness, distance to nearest settlement, dCOI, likelihood of human detection and time since wolf establishment. Because sample sizes allowed it, we added interactive terms between tree cover density and risk variables (distance to nearest settlement, dCOI, likelihood of human detection, and time since wolf establishment) to deepen our understanding of context-dependency in group size modulation. In order to account for baseline spatial and temporal variations, we added study areas as a random intercept in the models. We verified the performance and fit quality of each model by computing performance indices (*performance* R package^103^). We did not note any collinearity issues post-hoc (variance inflation factor < 2.5).

Across all models, explanatory variables were scaled and centred (x̄ = 0, sd = 1). We interpreted results based on statistical significance (*p* < 0.05) and effect sizes. All data processing and analyses were conducted in R software (v.4.6.0^104^).

## Acknowledgements

We thank the staff, rangers, students, and interns of Stelvio National Park and Fondazione Edmund Mach for their assistance with camera-trap deployments, checks and retrievals. We also thank Prof. Steeve Côté and Prof. Anne Loison for their reviews as external PhD examiners.

This work was financially supported by a PhD scholarship to CV co-funded by the Centre Agriculture Food Environment - C3A (University of Trento and the Fondazione Edmund Mach - FEM), and the Sustainable Development and Protected Areas Service of the Autonomous Province of Trento. CV, AC, FF, LL, FO, HCH, and FC were partially funded under the National Recovery and Resilience Plan (PNRR), Mission 4 Component 2 Investment 1.4 - Call for tender no. 3138 of 16 December 2021, rectified by Decree no. 3175 of 18 December 2021 of the Italian Ministry of University and Research funded by the European Union – NextGenerationEU (Project code CN_00000033, Concession decree no. 1034 of 17 June 2022 adopted by the Italian Ministry of University and Research, CUP D43C22001280006, J33C22001190001, B63C22000650007, and H43C22000530001, Project title ‘National Biodiversity Future Center – NBFC’). Views and opinions expressed are, however, those of the author(s) only and do not necessarily reflect those of the European Union or the European Commission, nor can they be held responsible for them.

## Author Contributions

CV, AC, and FC conceived the idea and designed the methodology, with contributions from LC and LP on the overarching study design; CV analysed the data and led the writing of the manuscript; CV, AC, VD, LL, FO, NS, LV collected and processed the data; LP, HCH, and FC supervised the work. All co-authors contributed critically to the drafts and all authors gave final approval for publication.

