## Supporting information for "Intra- and interspecific relationships of alpine ungulates reorganise along a wolf recolonisation gradient"

#### Details on the standardised camera-trap deployments

Each camera-trap was placed at the first accessible random location generated within each gridcell, excluding areas restricted by logistical issues such as safety concerns or private land access, and never placed within 500 meters of another camera from a neighbouring gridcell. At the selected site, the unbaited camera was positioned within 10-15 meters of the exact coordinates (accounting for GPS error) to maximise sighting probability, typically near game trails. The cameras were deployed about 50 centimeters above the ground, on a tree trunk or a wooden pole, facing north or perpendicular to the slope. Grass or tree branches for 2-3 m in front of the camera were cleared to reduce false triggers. To meet privacy laws and regulations, at each location, a signpost was placed to explain the purpose of the monitoring, and each camera-trap was locked. After deployment, the operator would record details such as time of the day, date, unique camera-trapping and SD card identifier codes, and trigger a few test images to make sure the camera-trap was working, but also to match the deployment time in case errors occurred during trapping. Camera-traps were Bushnell, Comitel, Cuddeback, ICU server, and Stealth Cam devices.

#### Details on the spatial covariates

To control for vegetational, topographic and anthropogenic effects on ungulate activity patterns, we extracted spatial covariates from four spatial raster layers, varying in resolution (from 5 to *ca* 150 meters).

##### Digital elevation model, slope and northness

We derived the digital elevation model (DEM) for the study area, namely the elevation in metres of the bare ground, from the TINITALY/1.1 dataset (Tarquini et al. 2023), which is available as a 10 m-cell size grid. From the DEM layer we then derived the slope, which is defined as the angle of inclination of the terrain, expressed in degrees, and the terrain aspect, representing the angle of absolute bearing of the overall terrain slope, with north considered as the origin (i.e., 0° for slopes facing north, 90° east, 180° south, and 270° west), using the corresponding functions in QGIS 3.28.6-Firenze (QGIS.org 2024). Northness was then calculated as the cosine of the aspect angle after converting it into radians, ranging from +1 (north-facing) to -1 (south-facing).

##### Tree cover density (TCD)

We derived a vegetation layer from high-resolution satellite data (Sentinel-2A/2B) provided by the European Environment Agency (EEA; https://doi.org/10.2909/486f77da-d605-423e-93a9-680760ab6791) under the EU Copernicus program. This raster provides information on proportional tree crown coverage per pixel at a 10 meters spatial resolution for 2018, with values ranging from 0 (completely non-tree-covered areas) to 1 (full tree cover).

##### Distance to nearest settlement

We extracted each land use type from a very high-resolution land use map (5 meters) of the study area, covering the years 2015 to 2020 (Marsoner et al. 2023). Specifically, we extracted human settlements (classes 11000, 11100, 11200, 11300, 11400, and 12100) and then generated a proximity raster by calculating the distance from each raster cell to the nearest feature, using the corresponding functions in QGIS 3.28.6-Firenze (QGIS.org 2024). The extraction area was defined as the bounding box of the four sampling grids, with a buffer of 10 km to reduce the edge effect.

##### Density of Cumulated Outdoor activity Index

We derived a layer depicting human mobility volume in the outdoors (density of Cumulated Outdoor activity Index, dCOI) following (Corradini et al. 2021). Specifically, the index is based on the crowdsourced Global Heatmap provided by STRAVA (San Francisco, CA, USA), which visualizes the outdoor activity tracks recorded and marked as public by users over the previous year (data extraction in October 2024). Specifically, for this study, we extracted data on all outdoor activities, primarily hiking, running, and cycling, at a zoom level 10 (i.e., 152 m pixel resolution). We then summed COI values within a 3×3 pixel moving window to account for the spatial context around each pixel and reduce potential pixel blindness, and we derived the dCOI. Further details on the representativeness of STRAVA data in both space and time are described by Thorsen et al. (2022) and Venter et al. (2023).

#### Detailed model outputs

***Table S1. Detection parameter estimates*** *(on the logit-scale) from the multi-species occupancy co-occurrence model. Estimates, standard errors (SE), z-values and associated p-values are provided per species and parameter. Significant effects are shown in bold.*

| Species | Parameter | | *Estimate* | *SE* | *z* | *p* |
|---|---|---|---|---|---|---|
| **Red deer** | **(Intercept)** | | **-0.98** | **0.06** | **-15.69** | **< 0.001** |
| | **Tree Cover Density** | | **-0.08** | **0.03** | **-3.23** | **0.001** |
| | Northness | | 0.04 | 0.02 | 1.63 | 0.104 |
| | **Slope** | | **-0.10** | **0.02** | **-4.45** | **< 0.001** |
| | **Camera-trap model** | Bushnell | **0.24** | **0.07** | **3.24** | **0.001** |
| | | Comitel | **0.52** | **0.07** | **7.05** | **< 0.001** |
| | | Cuddeback | **0.69** | **0.26** | **2.61** | **0.009** |
| | | ICU server | -0.09 | 0.10 | -0.88 | 0.381 |
| | | Stealth Cam | **-0.63** | **0.17** | **-3.70** | **< 0.001** |
| **Roe deer** | **(Intercept)** | | **-1.21** | **0.07** | **-17.19** | **< 0.001** |
| | **Tree Cover Density** | | **0.21** | **0.04** | **5.33** | **< 0.001** |
| | **Northness** | | **0.07** | **0.03** | **2.55** | **0.011** |
| | **Slope** | | **-0.09** | **0.03** | **-3.09** | **0.002** |

| | | | | | | |
|---|---|---|---|---|---|---|
| | **Camera-trap model** | Bushnell | -0.03 | 0.08 | -0.35 | 0.723 |
| | | Comitel | -0.08 | 0.09 | -0.89 | 0.372 |
| | | Cuddeback | -1.52 | 1.14 | -1.34 | 0.179 |
| | | ICU server | **-0.36** | **0.10** | **-3.47** | **< 0.001** |
| | | Stealth Cam | **-0.52** | **0.16** | **-3.32** | **< 0.001** |
| **Chamois** | **(Intercept)** | | **-2.12** | **0.14** | **-15.07** | **< 0.001** |
| | **Tree Cover Density** | | **-0.12** | **0.06** | **-2.17** | **0.030** |
| | Northness | | 0.01 | 0.04 | 0.25 | 0.805 |
| | **Slope** | | **0.13** | **0.05** | **2.80** | **0.005** |
| | Camera-trap model | Bushnell | 0.20 | 0.17 | 1.22 | 0.223 |
| | | Comitel | 0.33 | 0.17 | 1.93 | 0.054 |
| | | Cuddeback | -0.21 | 0.53 | -0.39 | 0.697 |
| | | ICU server | -0.52 | 0.49 | -1.06 | 0.291 |
| | | Stealth Cam | -0.55 | 0.40 | -1.36 | 0.172 |

**Table S2. Interspecific co-occurrence.** Occupancy parameter estimates (on the logit-scale) from the multi-species occupancy co-occurrence model. Estimates, standard errors (SE), z-values, and associated p-values are provided per species, species interaction and parameter. Significant effects are shown in bold.

| Species or Co-occurrence | Parameter | *Estimate* | *SE* | *z* | *p* |
|---|---|---|---|---|---|
| **Red deer** | **(Intercept)** | **1.57** | **0.39** | **4.07** | **< 0.001** |
| | Tree Cover Density | 0.13 | 0.35 | 0.36 | 0.718 |
| | Northness | 0.12 | 0.32 | 0.40 | 0.693 |
| | Slope | 0.57 | 0.31 | 1.85 | 0.065 |
| | Distance to nearest human settlement | -0.05 | 0.33 | -0.14 | 0.888 |
| | **density Cumulated Outdoor activity Index** | **-0.66** | **0.30** | **-2.25** | **0.025** |
| | Likelihood of human detection | -0.43 | 0.36 | -1.19 | 0.232 |
| | Time since wolf establishment | 0.32 | 0.32 | 1.00 | 0.318 |
| **Roe deer** | (Intercept) | 0.03 | 0.48 | 0.07 | 0.945 |
| | **Tree Cover Density** | **1.29** | **0.41** | **3.16** | **0.002** |
| | Northness | -0.03 | 0.39 | -0.09 | 0.929 |
| | Slope | 0.40 | 0.34 | 1.15 | 0.250 |
| | **Distance to nearest human settlement** | **-1.04** | **0.43** | **-2.42** | **0.016** |
| | density Cumulated Outdoor activity Index | -0.30 | 0.30 | -1.02 | 0.308 |
| | Likelihood of human detection | 0.07 | 0.31 | 0.23 | 0.815 |
| | **Time since wolf establishment** | **0.72** | **0.36** | **2.03** | **0.043** |
| **Chamois** | **(Intercept)** | **-1.02** | **0.51** | **-2.01** | **0.045** |
| | Tree Cover Density | -0.91 | 0.47 | -1.94 | 0.052 |
| | Northness | -0.57 | 0.42 | -1.36 | 0.174 |
| | **Slope** | **0.84** | **0.39** | **2.16** | **0.031** |
| | Distance to nearest human settlement | 0.11 | 0.38 | 0.28 | 0.782 |
| | density Cumulated Outdoor activity Index | -0.32 | 0.35 | -0.90 | 0.369 |
| | Likelihood of human detection | 0.27 | 0.36 | 0.73 | 0.463 |
| | Time since wolf establishment | -0.64 | 0.39 | -1.67 | 0.094 |
| **Red deer : Roe deer** | **(Intercept)** | **1.15** | **0.50** | **2.31** | **0.021** |
| | Tree Cover Density | -0.34 | 0.43 | -0.79 | 0.429 |
| | Northness | 0.29 | 0.41 | 0.71 | 0.476 |
| | Slope | -0.65 | 0.35 | -1.84 | 0.066 |
| | Distance to nearest human settlement | 0.57 | 0.43 | 1.32 | 0.188 |
| | density Cumulated Outdoor activity Index | 0.47 | 0.34 | 1.40 | 0.161 |
| | Likelihood of human detection | 0.50 | 0.37 | 1.36 | 0.174 |
| | **Time since wolf establishment** | **-0.92** | **0.37** | **-2.46** | **0.014** |
| **Red deer : Chamois** | **(Intercept)** | **1.07** | **0.49** | **2.16** | **0.031** |
| | Tree Cover Density | 0.50 | 0.46 | 1.09 | 0.275 |
| | Northness | 0.64 | 0.44 | 1.46 | 0.145 |
| | **Slope** | **-0.85** | **0.40** | **-2.12** | **0.034** |
| | Distance to nearest human settlement | -0.01 | 0.40 | -0.03 | 0.974 |

| | | | | | |
|---|---|---|---|---|---|
| | density Cumulated Outdoor activity Index | 0.05 | 0.39 | 0.14 | 0.890 |
| | Likelihood of human detection | -0.26 | 0.31 | -0.85 | 0.395 |
| | Time since wolf establishment | 0.29 | 0.40 | 0.72 | 0.474 |
| **Roe deer : Chamois** | **(Intercept)** | **-0.76** | **0.31** | **-2.43** | **0.015** |
| | Tree Cover Density | 0.24 | 0.30 | 0.79 | 0.427 |
| | Northness | -0.46 | 0.25 | -1.86 | 0.062 |
| | Slope | 0.48 | 0.25 | 1.90 | 0.058 |
| | Distance to nearest human settlement | 0.25 | 0.28 | 0.88 | 0.379 |
| | density Cumulated Outdoor activity Index | -0.43 | 0.32 | -1.37 | 0.169 |
| | Likelihood of human detection | -0.00 | 0.37 | -0.01 | 0.990 |
| | Time since wolf establishment | 0.03 | 0.27 | 0.12 | 0.908 |

**Table S3. Intraspecific grouping.** Model estimates, standard errors (SE), z-values, and associated p-values for ungulate group size as a function of environmental, predator, and anthropogenic variables. Red deer, roe deer, and chamois were modeled separately. P-values of single terms involved in significant interactions (-) were not shown due to little to no meaning in our models. This model was based on 2607 independent observations, and the fixed terms explained 0.4% of the total variance.

| Species | Predictors | Group size | | | |
|---|---|---|---|---|---|
| | | ***Estimate*** | ***SE*** | ***z*** | ***p*** |
| **Red deer** | **(Intercept)** | **0.53** | **0.05** | **9.74** | **<0.001** |
| | Time since wolf establishment (timeWolf) | 0.15 | 0.03 | 5.56 | - |
| | Tree cover density (TCD) | -0.13 | 0.01 | -10.40 | - |
| | Northness | -0.01 | 0.01 | -0.92 | 0.356 |
| | **Slope** | **-0.06** | **0.01** | **-4.96** | **<0.001** |
| | **Distance to nearest human settlement (distHumSett)** | **-0.06** | **0.01** | **-4.50** | **<0.001** |
| | density Cumulated Outdoor activity Index (dCOI) | -0.03 | 0.01 | -2.71 | - |
| | **TCD × timeWolf** | **-0.08** | **0.01** | **-7.27** | **<0.001** |
| | TCD × distHumSett | -0.00 | 0.01 | -0.35 | 0.729 |
| | **TCD × dCOI** | **0.09** | **0.01** | **7.70** | **<0.001** |
| **Chamois** | **(Intercept)** | **0.32** | **0.06** | **5.26** | **<0.001** |
| | Time since wolf establishment (timeWolf) | -0.03 | 0.07 | -0.51 | 0.610 |
| | Tree cover density (TCD) | 0.02 | 0.04 | 0.53 | - |
| | Northness | -0.02 | 0.03 | -0.62 | 0.536 |
| | **Slope** | **-0.12** | **0.03** | **-3.60** | **<0.001** |
| | Distance to nearest human settlement (distHumSett) | 0.00 | 0.04 | 0.09 | - |
| | density Cumulated Outdoor activity Index (dCOI) | -0.08 | 0.04 | -2.28 | - |
| | TCD × timeWolf | 0.01 | 0.03 | 0.35 | 0.729 |
| | **TCD × distHumSett** | **-0.07** | **0.04** | **-1.96** | **0.049** |
| | **TCD × dCOI** | **-0.11** | **0.04** | **-2.95** | **0.003** |
| **Roe deer** | **(Intercept)** | **0.16** | **0.02** | **8.47** | **<0.001** |
| | Time since wolf establishment (timeWolf) | 0.04 | 0.02 | 1.64 | 0.101 |
| | Tree cover density (TCD) | -0.01 | 0.02 | -0.54 | 0.588 |
| | Northness | -0.02 | 0.02 | -1.10 | 0.271 |
| | Slope | 0.02 | 0.02 | 1.11 | 0.269 |
| | Distance to nearest human settlement (distHumSett) | -0.01 | 0.02 | -0.28 | 0.782 |
| | density Cumulated Outdoor activity Index (dCOI) | 0.02 | 0.02 | 0.99 | 0.321 |
| | TCD × timeWolf | 0.03 | 0.02 | 1.57 | 0.117 |
| | TCD × distHumSett | 0.01 | 0.02 | 0.72 | 0.473 |
| | TCD × dCOI | 0.03 | 0.02 | 1.14 | 0.256 |

#### Visualisation of the overall neutral co-occurrence among red deer, roe deer, and chamois

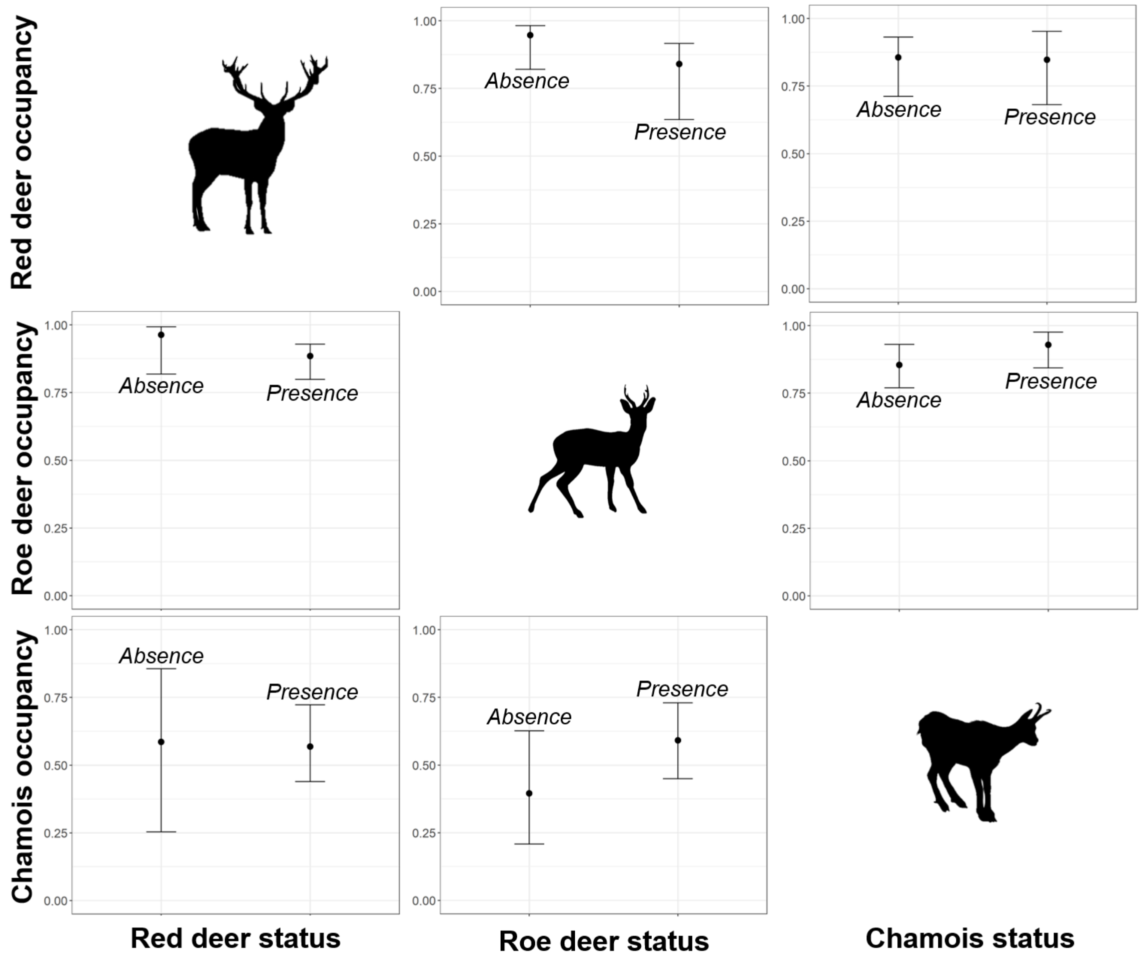


***Figure S1.*** *Occupancy of red deer (**top**), roe deer (**middle**) and chamois (**bottom**) in function of the presence or absence of red deer (**left**), roe deer (**middle**) and chamois (**right**). Dots show mean probability of occupancy and lines the 95% confidence intervals. Occupancy probabilities were estimated by setting all covariates to their mean values.*

### Visualisation of the time-to-independence thresholds

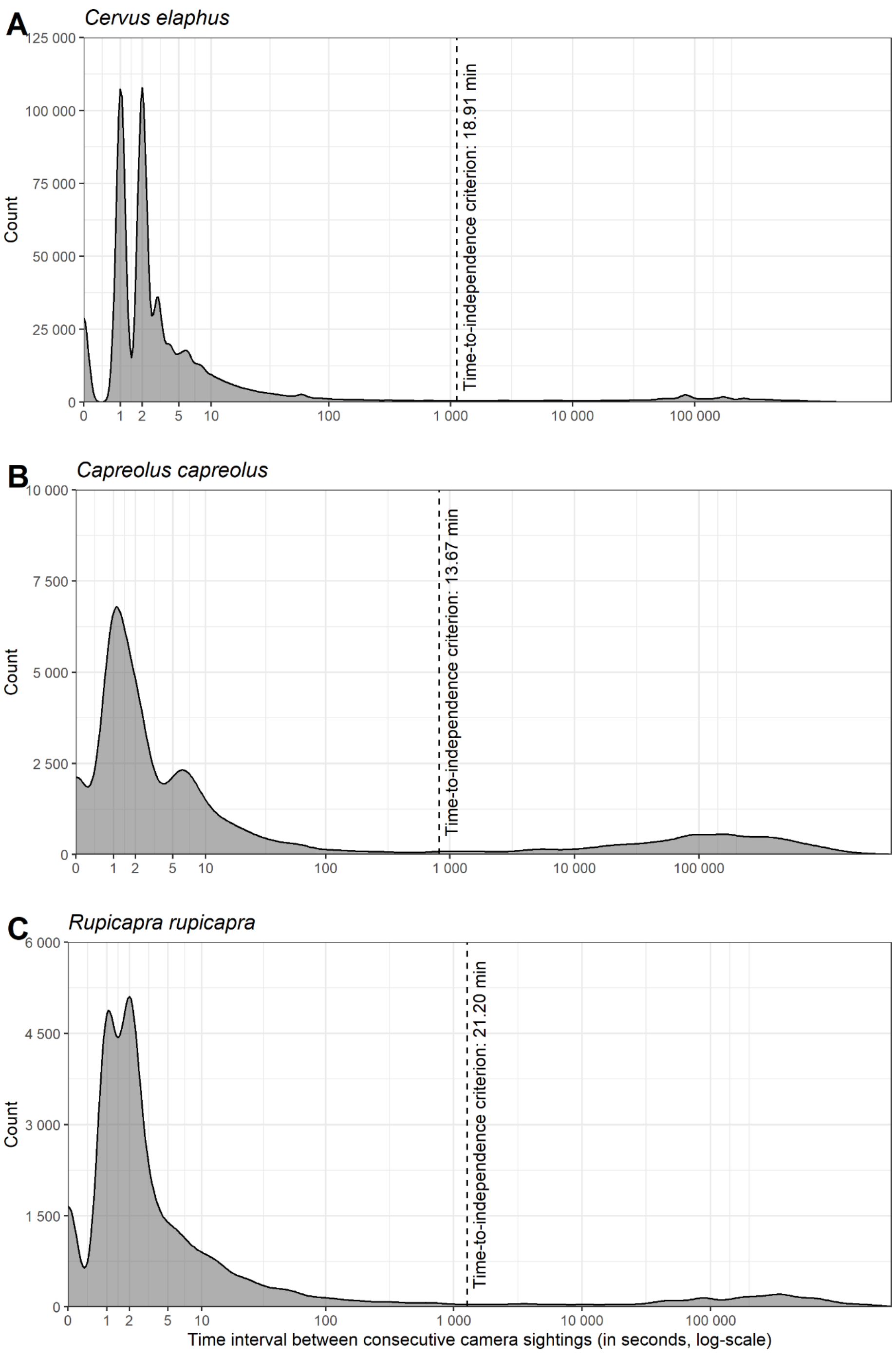


***Figure S2.*** *Distribution of time intervals (in seconds, transformed with a logarithmic scale) between consecutive camera sightings of (**A**) red deer, (**B**) roe deer, and (**C**) chamois. The vertical dashed lines show the species-specific time-to-independence thresholds, determined using the bout-ending criterion (Luque & Guinet 2007) by fitting a mixture of two Poisson processes to the time intervals between consecutive camera sightings.*
